# ProteoMixture: A Cell Type Deconvolution Tool for Bulk Tissue Proteomics Data

**DOI:** 10.1101/2023.07.27.550810

**Authors:** Pang-ning Teng, Joshua P. Schaaf, Tamara Abulez, Brian L. Hood, Katlin N. Wilson, Tracy J. Litzi, David Mitchell, Kelly A. Conrads, Allison L. Hunt, Victoria Olowu, Julie Oliver, Fred S. Park, Marshé Edwards, AiChun Chiang, Praveen-Kumar Raj-Kumar, Christopher M. Tarney, Kathleen M. Darcy, Neil T. Phippen, G. Larry Maxwell, Thomas P. Conrads, Nicholas W. Bateman

**Author notes:** Authors contributed equally (Pang-ning Teng, Joshua P. Schaaf). Corresponding Authors: Nicholas W. Bateman, Women’s Health Integrated Research Center at Inova Health System, 3289 Woodburn Road, Suite 370, Annandale, VA 22003; and Thomas P. Conrads, Women’s Health Integrated Research Center at Inova Health System, 3289 Woodburn Road, Suite 370, Annandale, VA 22003.

## Abstract

Numerous multi-omic investigations of cancer tissue have documented varying and poor pairwise transcript:protein quantitative correlations and most deconvolution tools aiming to predict cell type proportions (cell admixture) have been developed and credentialed using transcript-level data alone. To estimate cell admixture using protein abundance data, we analyzed proteome and transcriptome data generated from contrived admixtures of tumor, stroma, and immune cell models or those selectively harvested from the tissue microenvironment by laser microdissection from high grade serous ovarian cancer (HGSOC) tumors. Co-quantified transcripts and proteins performed similarly to estimate stroma and immune cell admixture in two commonly used deconvolution algorithms, ESTIMATE and Consensus^TME^ (r ≥ 0.63). Here we have developed and optimized protein-based signatures to estimate cell admixture proportions and benchmarked these using bulk tumor proteomics data from over 150 HGSOC patients. The optimized protein signatures supporting cell type proportion estimates from bulk tissue proteomics data are available at (https://lmdomics.org/ProteoMixture/.

## Introduction

The ovarian cancer tumor microenvironment (TME) includes various cell types such as tumor, stroma, and immune cells that can regulate tumor development and progression (1, 2). Immune cell populations, including tumor-associated/infiltrated lymphocytes (TILs), in the TME have been shown to impact cancer prognosis and response to neoadjuvant chemotherapy (NACT) (3, 4). Proteogenomic analyses of high grade serous ovarian cancer (HGSOC) to date have largely utilized bulk tumor collections that contain widely varying admixtures of diverse cell types (5, 6). Our group (7) and others (8) have shown that variations in the proportions of different cellular populations within the TME can impact correlation with different HGSOC prognostic molecular subtypes (9–13). Improved characterization of cell admixture contributions to the bulk tissue proteome will support refinement of proteogenomic signatures from bulk and enriched cell type collections.

Deconvolution of cell type proportions (cell admixture) from bulk expression data has previously been achieved by quantifying the enrichment of cell type-associated gene expression signatures. Current deconvolution tools include Estimation of STromal and Immune cells in MAlignant Tumor tissues using Expression data (ESTIMATE) (14), xCell (15), Microenvironment Cell Populations-counter (MCP-counter) (16) and CIBERSORTx (14–18). Some of these tools have also recently been merged into an integrated tool, Consensus^TME^ (18), enabling prediction of cell admixture for 18 different cell types including fibroblasts, endothelial cells, and 16 immune-related cell types. These tools have been developed using transcript-level data, which exhibits limited correlation to proteome abundances in cancer cells and tissues including HGSOC (6, 19), largely due to translational regulation (20). Thus, there remains a paucity of data investigating the applicability of these signatures for characterizing cellular admixture within proteome data in HGSOC tissues. Very recent efforts by Feng et al. (21) described the Decomprolute tool which enables prediction of immune cell signatures using proteomic data and established deconvolution tools across various organ site malignancies, including ovarian cancer. Motivated by this work as well as recent efforts by our group correlating stromal cell admixture with prediction of the mesenchymal (MES) subtype (7), a molecular subtype correlating with poor disease prognosis in HGSOC, we examined proteomic signatures of tumor, stroma, and immune cell admixture in HGSOC.

Our study describes an evaluation of the performance of matched transcriptome and proteome data generated from a contrived admixture series of HGSOC tumor, stroma/fibroblasts, and immune cells using existing deconvolution and prognostic molecular subtype prediction tools. We have further investigated the impact of cell type admixture on the correlation of protein and transcript abundances. We describe optimized protein signatures for tumor, stroma, and immune cell admixtures and their performance in classifying proteome data from enriched and bulk tissue collections for multiple, independent HGSOC patient cohorts. We provide these signatures as part of a publicly available tool, ProteoMixture, supporting cell type deconvolution from bulk tissue proteomic data (https://lmdomics.org/ProteoMixture/).

## Results

### Proteogenomic Analysis of HGSOC Cell Admixture Models

We generated cell admixtures consisting of defined percentages of tumor, stroma, and immune cell populations from either cultured cell line models or laser microdissection (LMD) enriched tissues. Specifically, *in vitro* cell line admixtures were generated using HGSOC tumor cells (OVCAR-3) (22), fibroblast cells (to mimic stromal cells) established from an *in situ* ovarian cancer (23, 24), and a model of T-cells (Jurkat) (25). LMD harvested tissue mixtures were generated using enriched populations of tumor, stroma, and immune-infiltrated stroma cells pooled from five women diagnosed with HGSOC (Figure 1, Supplementary Table 1-5). Global proteome and transcriptome analyses of cell admixtures were performed using a quantitative, multiplexed proteomic approach employing tandem mass tags (TMT) and liquid chromatography, high-resolution tandem mass spectrometry (LC-MS/MS), and RNA sequencing (RNA-seq), respectively. Global proteome and transcriptome analyses quantified 6,683 ± 783 proteins and >20,000 transcripts across all admixtures (Supplementary Tables 6-9).

**Figure 1:**
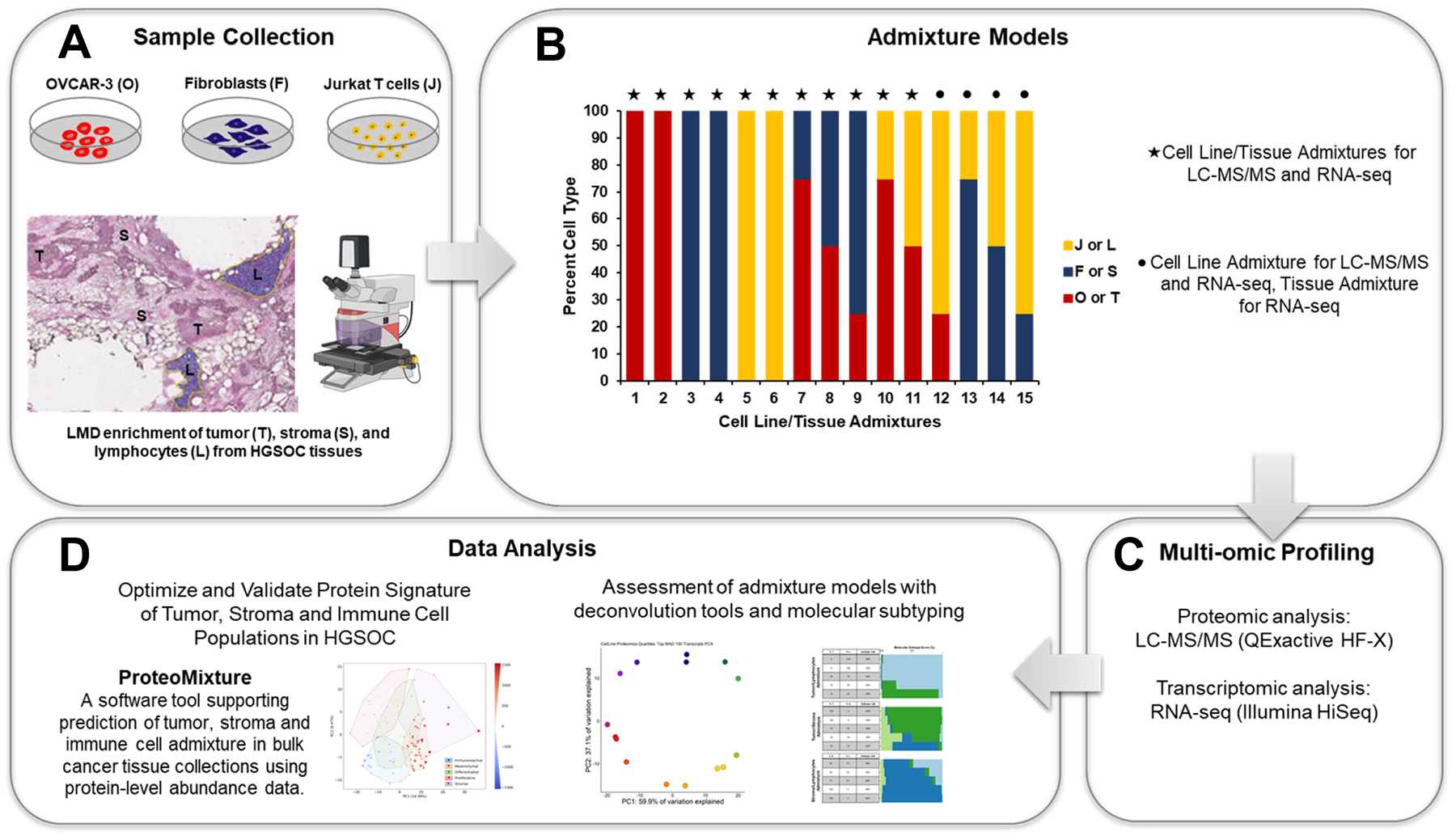
Analytical Workflow Supporting Proteogenomic Analysis of Cellular Admixture in High Grade Serous Ovarian Cancer (HGSOC). A & B: Cell admixtures were generated by mixing varying amounts cell type-associated peptide digests or extracted RNA collected from cell line models and subpopulations of cell of interest from HGSOC tissues. C & D: Cell admixtures were analyzed by LC-MSMS and RNA-seq followed by assessment in established deconvolution tools and prognostic molecular subtype prediction tools. Protein signatures predicting tumor, stroma, and immune cell admixture were optimized and validated in independent HGSOC cohorts. Protein-level signatures were distilled into a publicly available tool supporting cell type deconvolution from bulk tissue proteomic data (https://lmdomics.org/ProteoMixture/).

Proteome and transcriptome data from cell admixtures reflecting quartile percentages of tumor, stroma, and immune cell types were evaluated by principal component analysis (PCA) (Figure 2). PCA of the top 100 variably abundant proteins and transcripts by median absolute deviation (MAD) showed that admixtures that comprise one predominant cell type (tumor, stroma, or immune cells) form largely distinct clusters that transition to clusters of related sample compositions across cell type dilution series (Figure 2). PCA analysis of the top 100 variably abundant proteins explained 56.6 ± 4.67% and 41.45 ± 6.15% (≤ 14.84% CV, Figure 2A and 2B) of the variance between cell admixture conditions generated using cell lines or tissue samples, respectively. Similar analyses of the top 100 variably abundant transcript explained 74.30 ± 10.32% and 23.85 ± 8.98% (≤ 37.65% CV, Figure 2C and 2D) of the variance between cell admixtures generated using cell lines or tissue samples. Correlation analysis of 6,097 proteins and transcripts co-quantified across non-admixed cell populations of interest identified that purified tumor (Spearman’s ρ = 0.48 ± 0.005) and immune enriched samples (Spearman’s ρ = 0.483 ± 0.002) exhibited the highest mean correlation of transcript and protein abundances while enriched stroma samples were significantly lower (Spearman’s ρ = 0.32 ± 0.04, Supplementary Figure 1A, Supplementary Table 10). When we examined tumors admixed with other cell types, the correlation between transcript and protein abundances decreased as the percentage of stroma or lymphocytes increased (Supplementary Figure 1B and 1C, Supplementary Table 10).

**Figure 2:**
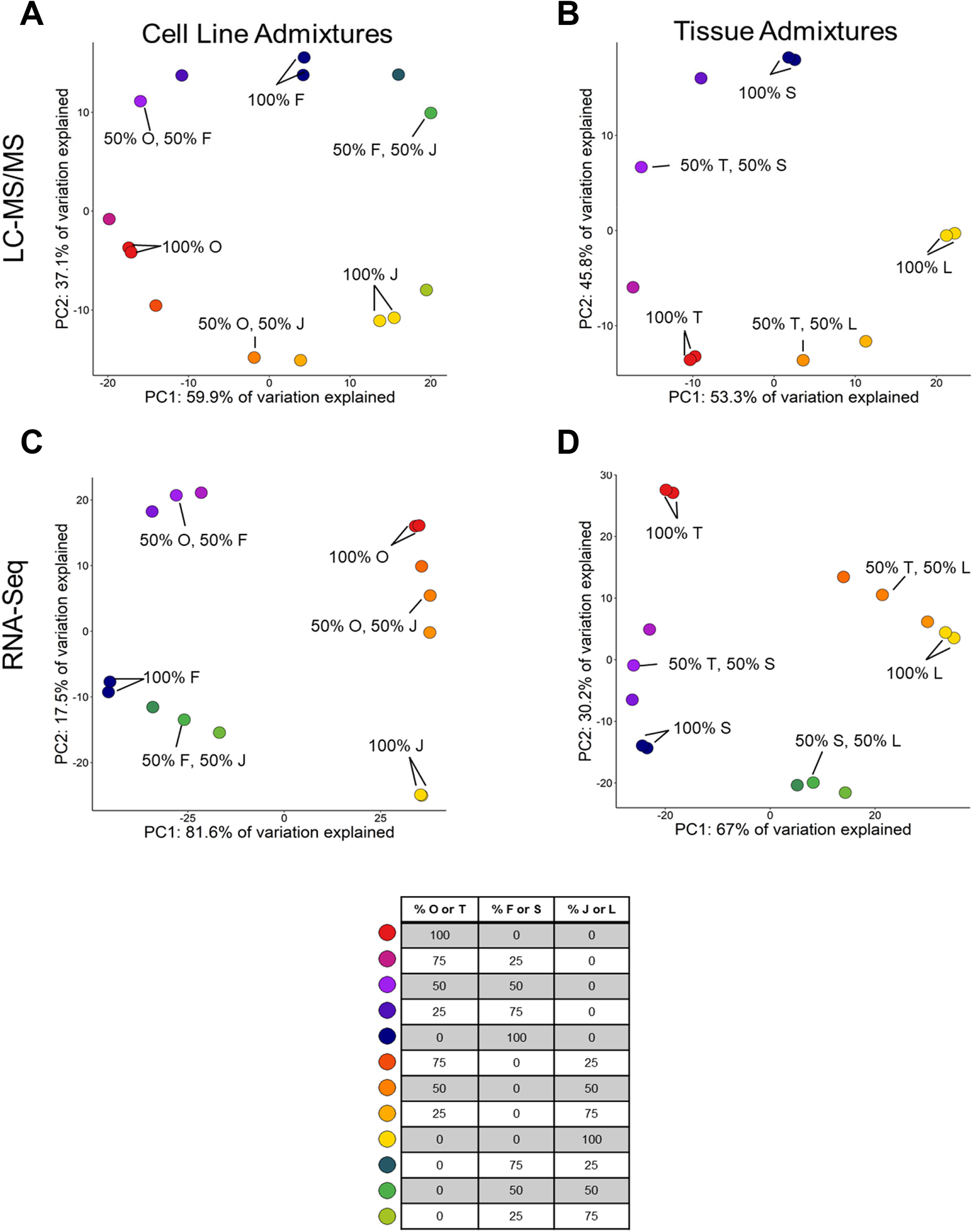
Principal Component Analysis (PCA) of Cell Admixture Models. PCA analyses using the top 100 most variably abundant proteins (median absolute deviation) in cell line admixtures (A) and LMD enriched cell populations from HGSOC tissues (B). PCA analyses using the top 100 most variably abundant transcripts in cell line admixtures (C) and LMD enriched cell populations from HGSOC tissues (D). Abbreviations: OVCAR-3 (O), Fibroblast (F), Jurkat T cells (J), Tumor (T), Stroma (S), Lymphocytes (L).

### Performance of Cell Admixture Models using Established Deconvolution and Molecular Subtype Classification Tools

We analyzed proteome and transcriptome data with previously published cell deconvolution tools established using gene expression or transcript-level data, i.e., ESTIMATE (14) and Consensus^TME^ (18) (Supplementary Tables 11-18). We first assessed the overlap between protein and transcript features comparing 1) stroma and immune score signature gene candidates in ESTIMATE, and 2) fibroblast and immune score gene signature candidates in Consensus^TME^. An average of 46% of gene signature constituents across these tools were co-quantified in both global proteome and transcriptome datasets (Supplementary Figure 2A). We then correlated resulting ESTIMATE and Consensus^TME^ scores with proportional cell population admixtures for cellular subpopulations of interest (Figure 3A-D). The ESTIMATE stromal score (Pearson’s r > 0.9, *p value* < 0.05) and the Consensus^TME^ fibroblast score (Pearson’s r > 0.9, *p value* < 0.05) were positively correlated with the percent stroma cell admixture from HGSOC tissues using proteomic or transcriptomic data (Figure 3A and B, Supplementary Table 19). In cell line admixtures, higher correlations between Consensus^TME^ fibroblast score and percent fibroblast (Pearson’s r > 0.88, *p value* < 0.05) were observed as compared to correlation between ESTIMATE stromal score and percent fibroblast (Pearson’s r > 0.82, *p value* > 0.05) (Figure 3, Supplementary Table 19). Immune scores from ESTIMATE or Consensus^TME^ scaled proportionally with increasing immune cell populations using co-quantified proteins (Pearson’s r > 0.9, *p value* < 0.05) from cell line admixtures or transcripts (Pearson’s r > 0.9, *p value* < 0.05) from HGSOC tissue admixtures as input (Figure 3C and D, Supplementary Table 19). ESTIMATE tumor purity scores positively correlated with percent OVCAR-3 and tumor cell collections between proteomic and transcriptomic data (Pearson’s r > 0.9, *p value* < 0.05) (Supplementary Figure 2B, Supplementary Table 19). Global proteome data collected from LMD enriched HGSOC tissue admixtures was also analyzed using a proteome-based immune cell deconvolution tool, Decomprolute (21). Decomprolute scores for CD8+ T cells and B cells were highly correlated with the percent immune cell HGSOC tissue admixture conditions (Pearson’s r > 0.94, *p value* < 0.01) (Supplementary Figure 3). Decomprolute also characterized LMD enriched lymphocyte cell populations as being comprised of predominantly CD8+ T cells (Decomprolute score = 0.42 ± 0.01) and B cells (Decomprolute score = 0.28 ± 0.02) (Supplementary Table 20).

**Figure 3:**
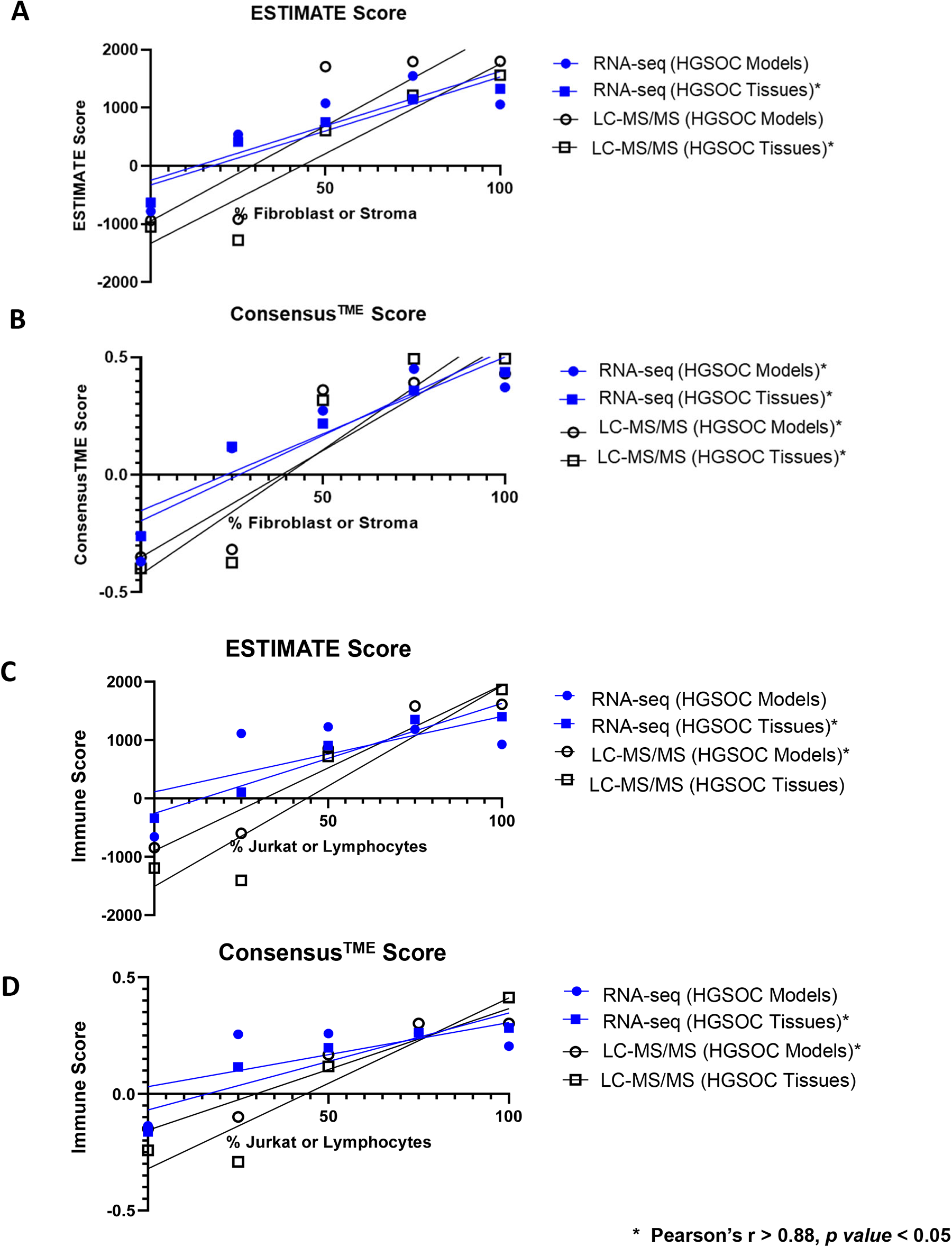
Comparison of Protein and Transcript-Level ESTIMATE and Consensus^TME^ Deconvolution Scores across Cell Admixture Conditions. A. Correlation analysis of transcriptomic (RNA-seq) and proteomic (LC-MS/MS) scores for quartile dilutions from cell lines and tissue collections corresponding to ESTIMATE stroma scores (A), Consensus^TME^ fibroblast scores (B), ESTIMATE immune scores (C), or Consensus^TME^ immune scores (D), (Supplementary Tables 11-19).

Prognostic molecular subtypes derived from gene signature analyses have been described in HGSOC patient tumors (10), from which the immunoreactive (IMR) and mesenchymal (MES) correlate with known immune or stroma cell populations, respectively (8, 12). We explored the impact of cell admixture using transcriptome data from LMD enriched tissue samples on prognostic molecular subtype classifications using the consensusOV tool (10). Cell admixtures predominated by tumor cells were largely classified as differentiated (DIF) subtype. Admixtures with high proportions of stroma or immune cells were classified as MES or IMR subtypes, respectively (Figure 4). Tumor and lymphocyte cell admixture classifications transitioned from IMR to DIF subtype between 20-50% lymphocytes. Tumor and stroma admixture classifications transitioned from DIF to MES between 50-75% stroma cells. Lastly, stroma and lymphocyte admixtures transitioned from MES to IMR between 50-75% lymphocytes.

**Figure 4:**
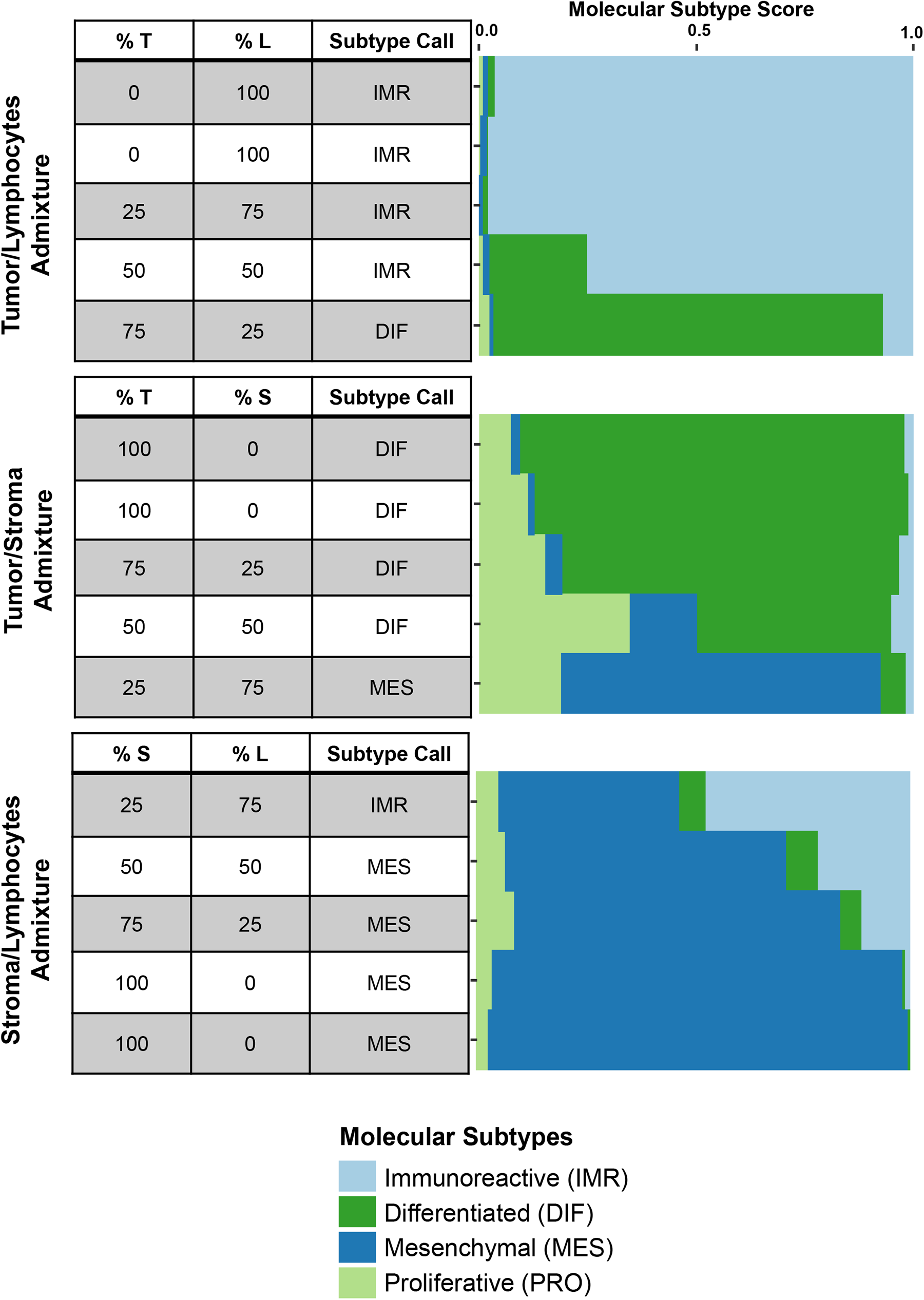
Comparison of Prognostic Molecular Subtypes in High Grade Serous Ovarian Cancer (HGSOC) across Cell Admixture Conditions. Transcriptome data from tumor/stroma, tumor/lymphocytes, and stroma/lymphocytes LMD enriched tissue admixtures were analyzed by consensusOV to classify molecular subtype. Data shows the percent admixture condition, top subtype call, and the molecular subtype scores for each cell admixture condition.

### Optimization of Protein Signatures Enabling Prediction of Tumor, Stroma, and Immune Cell Admixture in Proteomic data from Enriched and Bulk HGSOC Tissues

Cell type-associated protein signatures were generated from differential analysis of LMD enriched tumor, stroma, and immune cell-infiltrated populations (not admixed) from HGSOC tissues pooled from five patient tumors (Supplementary Table 1). More than 550 cell type-associated proteins were identified (Supplementary Figure 4, Supplementary Table 21). Recursive feature elimination (RFE) optimally selected 28 proteins uniquely elevated in tumor cells, 263 proteins elevated in stroma, and 268 proteins elevated in immune-enriched tissues collections (Figure 5A, Supplementary Table 21). We compared these cell type-associated protein signatures with known proteomic markers of LMD enriched HGSOC tumor cell populations (7), the gene signature candidates used in ESTIMATE and Consensus^TME^ deconvolution tools (14, 18), and with molecular subtype classification signatures (10, 12, 13) (Figure 5B). All 28 tumor signature proteins were unique relative to previously described feature sets, while 97 (37%) and 35 (13%) of the stroma and immune signature proteins, respectively, overlapped with marker transcripts from previously described signatures (Figure 5B, Supplementary Table 22). Transcript-protein correlation of proteins co-identified in cell type transcript signatures from deconvolution tools (ESTIMATE, Consensus^TME^) (14, 18), molecular subtype classification algorithms (12, 13), and protein signatures from LMD HGSOC (7) was significantly greater when compared to transcript-protein correlation of all protein signatures (*p value* < 0.0001) (Figure 5B and Supplementary Figure 5).

**Figure 5:**
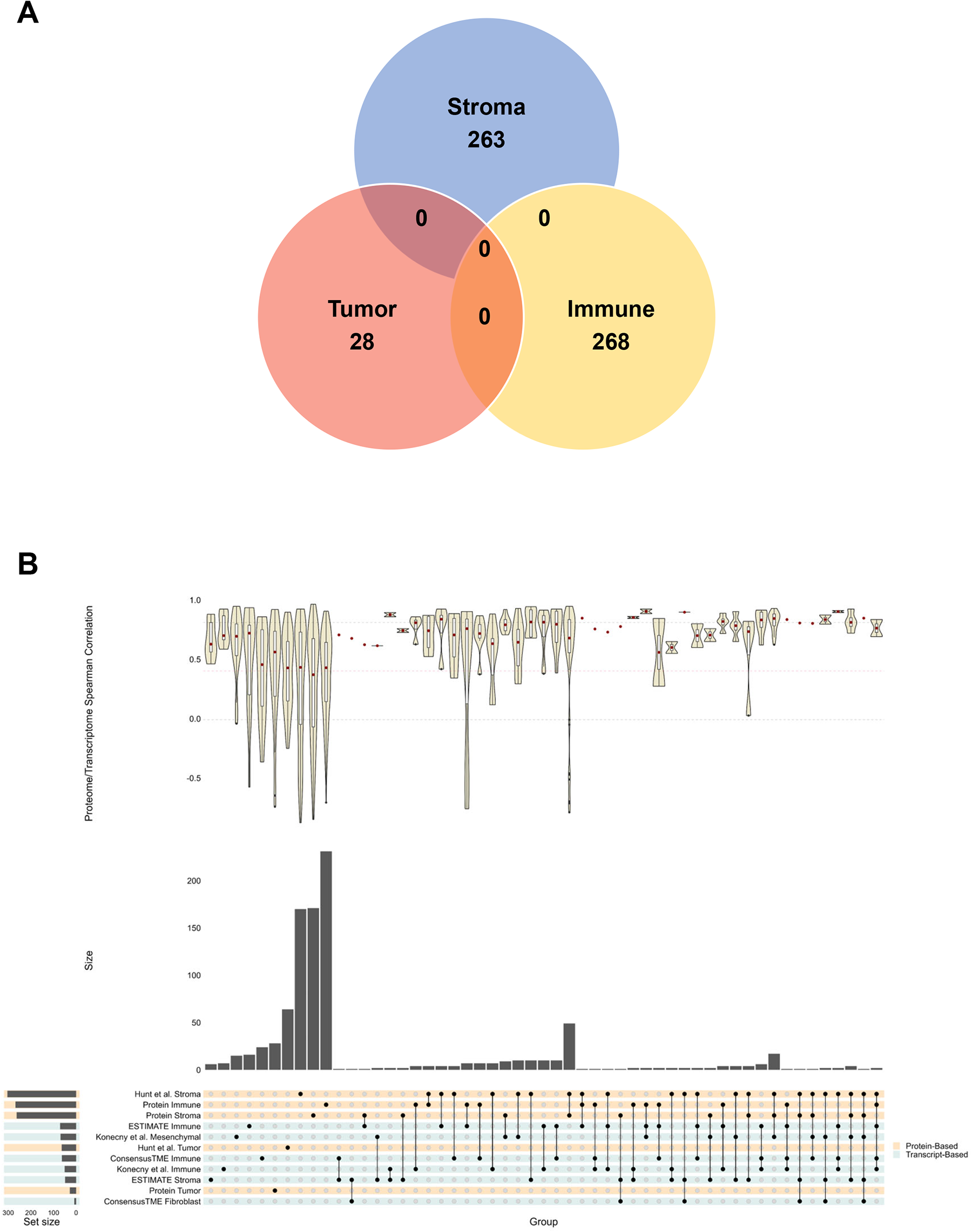
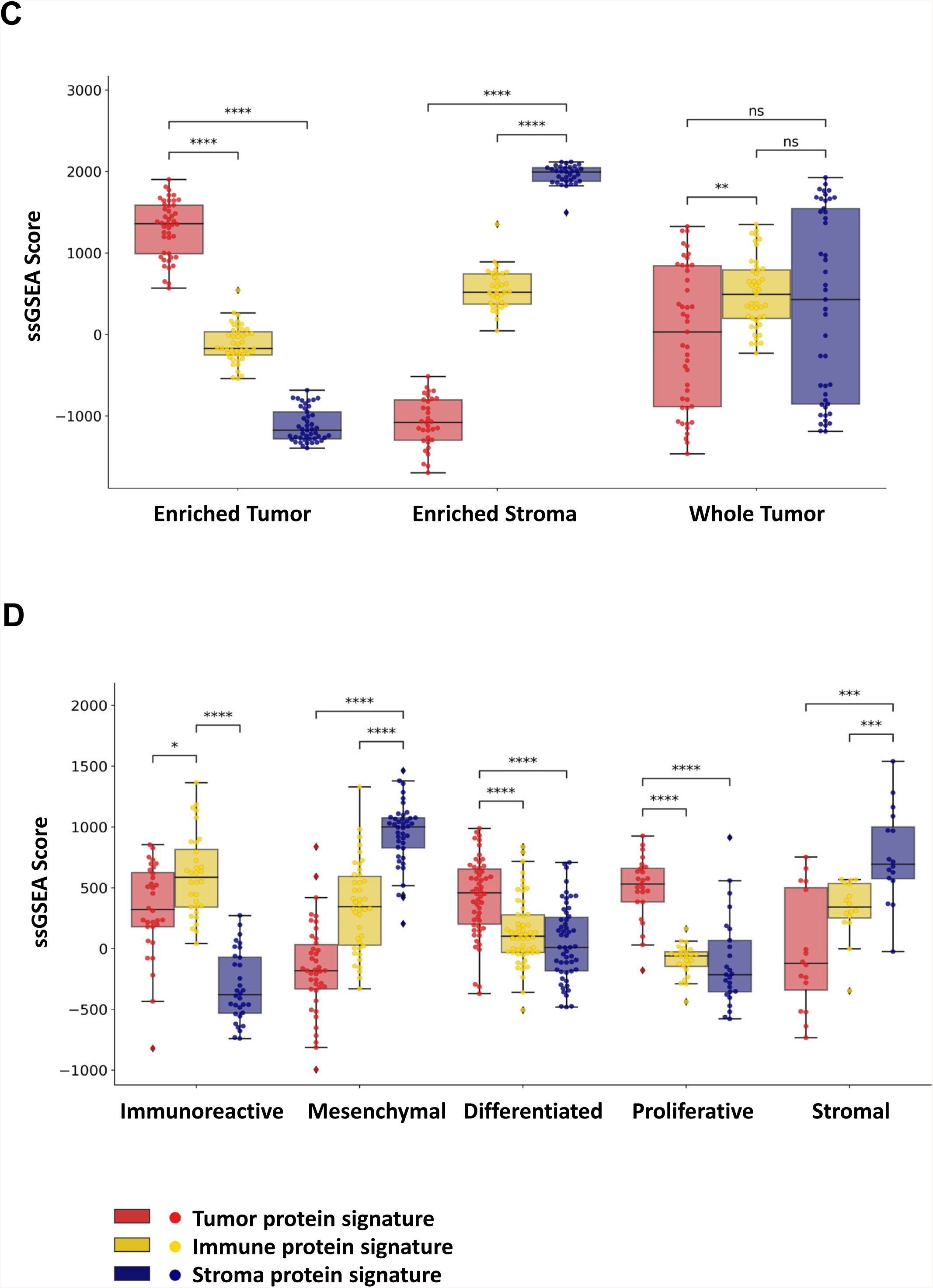
Optimization of Protein Signatures Enabling Prediction of Tumor, Stroma, and Immune Cell Admixture in Proteomic data from Enriched and Bulk HGSOC tissues. A: Venn diagram comparing optimized protein signature candidates prioritized for tumor, stroma and immune enriched cell populations. B: Integrated upset plot comparing optimized protein signatures with previously published cell type or molecular subtype protein and gene signatures with companion violin plots showing protein and transcript correlation distributions for overlapping features of interest; median correlation distributions denoted with red box. C: Assessment of protein signatures in proteomic data from enriched and bulk HGSOC tissues (Hunt et al. (7), n = 9 HGSOC patients); *0.01 < p ≤ 0.05, **p ≤ 0.001, ****p ≤ 0.0001 from Mann-Whitney Testing. D. Verification of protein signatures in proteomic data from bulk HGSOC tissues (Zhang et al. (6), n = 169 HGSOC patients); *0.01 < p ≤ 0.05, **p ≤ 0.001, ****p ≤ 0.0001 from Mann-Whitney Testing.

We assessed the predictive accuracy of our RFE-generated signatures for classifying tumor, stroma, and immune cells from LMD enriched tumor and stroma (7), and from bulk tissue collections from tumors of > 150 HGSOC patients (6, 7) (Figure 5C-D). As anticipated, the LMD enriched tumor exhibited significantly elevated tumor scores while enriched stroma exhibited significantly elevated stroma scores (Figure 5C). Bulk tissue collections exhibited more variable correlation with protein-derived tumor, stroma, and immune enriched signatures, consistent with the ≤ 56% tumor purities described for these samples (7). We also investigated the performance of our protein-level signatures in proteome data generated for bulk tissue collections recently described (6) by CPTAC for an additional independent cohort of >150 HGSOC tumors (Figure 5D). Tumors classified as DIF or proliferative (PRO) molecular subtypes by Zhang *et al.* exhibited the highest enrichment with the tumor protein signature. Further, tumors classified as MES or stromal had the highest single-sample gene set enrichment analysis (ssGSEA) scores with our stroma protein signature, while tumors classified as IMR scored highest with the immune protein signature. PCA analysis of the most variably abundant protein features (top 25% median absolute deviation) in the proteomic data revealed clustering of patient tumors by molecular subtype classification (Supplementary Figure 6). Overlaying the ssGSEA scores calculated for each patient tumor using our stroma protein signature (Supplementary Figure 6A), immune protein signature (Supplementary Figure 6B), and tumor protein signature (Supplementary Figure 6C) reveals that correlation with higher stroma (MES, stromal), immune (IMR) or tumor (DIF, PRO) molecular subtypes, respectively, as anticipated. We integrate these signatures into a publicly available tool supporting cell type deconvolution from bulk tissue proteomic data available here: https://lmdomics.org/ProteoMixture/.

## Discussion

Our study assessed matched proteome and transcriptome data generated from *in vitro* and *in situ*-derived admixtures of common HGSOC TME cell types using established transcript-based tools for cell type deconvolution and prognostic molecular subtype classification. Unique protein signatures were developed to classify tumor, stroma, and immune cell populations within admixture HGSOC tissue samples using protein-level data. Admixtures of tumor, stroma, and immune cell populations exhibit unique proteomic and transcriptomic profiles which drive sample clustering in unsupervised analysis. Our group showed previously that enrichment of tumor and stroma cells can result in markedly different proteome and transcriptome profiles for a given sample (7). Our findings agree with these results and further show that immune cell admixture can contribute unique proteogenomic abundance alterations, impacting the molecular profile of a given sample. We also investigated the impact of cell admixture on the abundance of co-quantified proteins and transcripts, as we previously observed that enriched stroma exhibits lower correlation trends of these features compared with enriched tumor cell populations (7). We identified median correlation distributions for proteins and transcripts in enriched tumor (∼0.48) and stromal cell (∼0.32) populations in this study, consistent with trends previously observed in HGSOC (7, 26). We further identified that correlation of protein and transcripts co-quantified in immune admixed conditions exhibit comparable median correlation trends as tumor cells (∼0.48), suggesting higher coordination of protein and transcript abundances is reflective of not just tumor cells but immune cell populations as well.

We also assessed the overlap of co-quantified protein and transcript features from *in vitro* or *in situ* sample collections with gene candidates from two previously described cell deconvolution tools, ESTIMATE (14) and Consensus^TME^ (18). Our efforts show that protein and transcript-level data exhibit comparable performance to predict immune and stroma/fibroblast cell admixture, including estimation of tumor purity. Although less than 50% of gene signature candidates for these tools were quantified at the protein-level, these features exhibited high quantitative correlation in transcript-level data (Supplementary Figure 5), likely explaining this comparable performance. A limitation of our study is that immune-enriched tissue samples were generated by collecting and pooling immune admixed cell populations within the tumor microenvironment of several HGSOC patient tumors, precluding our ability to characterize unique subpopulations of immune cells beyond generic assessments of immune cell admixture through assessment of companion “immune scores”. To this end, our analysis using Decomprolute (21) revealed high correlation of immune admixed conditions with CD8+ T cells and B cells from LMD enriched lymphocyte samples (Supplementary Figure 3). Future efforts will focus on generating protein-level signatures which enable classification of distinct immune cell populations involved in immune surveillance in HGSOC, such as CD8+ T cells, activated CD4+ T cells, and plasma cells (4). Other cell type deconvolution tools using proteomic data that have been developed includes scpDecov (27), which was developed based on single cell proteomic data, but we did not assess performance of this tool due to the limited feature sets quantified in single cell proteomic data in comparison with the multiplexed global proteomics workflow we applied in our study.

Using transcript-level data, we demonstrate that cell admixture directly impacts the classification of prognostic molecular subtypes (10) where we identify that admixture conditions consisting of >25% of a given cell population can impact the prediction of molecular subtype classification. Our findings resonate previous results showing that cellular heterogeneity directly impacts prognostic molecular subtype classifications in HGSOC (7). Molecular signatures generated using transcriptomic abundances from bulk tumor samples can lead to misinterpretation of their corresponding protein-level abundances due to stromal influences and the limited concordance between protein and transcript abundances (28), a finding we also observe in this study and in our prior work (7). These analyses underscore the importance of assessing tumor cellularity and sample heterogeneity when interpreting prognostic molecular subtype classifications for a given sample.

We prioritized protein signatures based on proteins uniquely elevated in tumor, stroma, and immune-enriched cell populations using recursive feature elimination. We then regressed these features relative to cell admixture dilution conditions to select an optimized set of proteins demonstrating strong performance to estimate tumor, stroma and immune cell admixture in protein-level data alone. Our tumor, stroma, and immune protein signatures were largely unique relative to gene signatures utilized in existing cellular deconvolution or molecular subtype classification tools (12, 14, 18). Further, most of our protein signature candidates exhibited lower correlation with cognate transcript abundances relative to features identified in previously described gene signatures. Our protein signatures successfully classified enriched tumor or stroma HGSOC samples using proteome data alone (7) and further correlated with molecular subtype classifications estimated from bulk HGSOC tissue collections (6), where tumors highly admixed with stromal cell populations (i.e., MES and stromal molecular subtypes) exhibit the highest stroma protein signature scores, tumors highly admixed with immune cell populations (i.e., IMR molecular subtype) exhibit high immune protein signature scores, and tumors exhibiting higher tumor purity (i.e., DIF and PRO molecular subtypes) exhibited high tumor protein signature scores.

To make these signatures readily accessible to the research community, we have developed ProteoMixture (https://lmdomics.org/ProteoMixture/), a tool for predicting cell types in proteomic data from HGSOC tissues. Cell type-unique protein signatures were validated and optimized for HGSOC, a disease typified by marked heterogeneity within the tumor microenvironment. Future efforts will be focused on assessing and refining protein signatures for characterizing cell admixtures in other organ sites beyond HGSOC. To this end, we investigated performance of global proteome data in the ProteoMixture tool from a recently described cohort of tumors collected from n=87 patients diagnosed with lung adenocarcinoma (29). We correlated ProteoMixture and ESTIMATE scores calculated from transcript-level data and identified high correlation between stroma (Spearman’s ρ = 0.699) and immune scores (Spearman’s ρ = 0.796) and significant, although lower correlations, between tumor purity estimates and tumor score (Spearman’s ρ = 0.47) (data not shown). These analyses demonstrate proof-of-concept of the utility of our protein signatures to assess tumor, stroma, and immune cell admixture in other organ site malignancies. The lower correlation of non-HGSOC tumor scores necessitates further selection and refinement using enriched cell populations from organ-site specific collections which will be the focus of future efforts.

## Materials and Methods

### Cell Culture

Human cell lines OVCAR-3 (NIH:OVCAR-3) and Jurkat T cells (Clone E6-1) were purchased from ATCC (Manassas, VA, USA). Human ovarian serous cancer associated fibroblasts (primary cells) were purchased from Vitro Biopharma, Inc. (Golden, CO, USA). OVCAR-3 was cultured in RPMI-1640 medium (ATCC) supplemented with 20% fetal bovine serum (FBS) (ATCC), Penicillin (100 IU/mL)-Streptomycin (100 µg/mL) (ATCC), and 0.01 mg/mL insulin (Sigma-Aldrich, St. Louis, MO, USA). Jurkat T cells were cultured in RPMI supplemented with 20% FBS (ATCC), Penicillin (100 IU/mL)-Streptomycin (100 µg/mL) (ATCC) and maintained at 1 x 10^5^ – 1 x 10^6^ cells/mL density. Ovarian serous cancer associated fibroblasts were cultured in VitroPlus III Low Serum Complete Medium (Vitro Biopharma, Inc.). All cells were maintained at 37 °C and 5% CO_2_. OVCAR-3 and ovarian serous cancer associated fibroblasts were grown in a monolayer on 10 cm^2^ tissue culture dishes (Fisher Scientific, Inc., Hampton, NH). At 70% confluency, with maintained healthy morphology, cells were washed twice with Dulbecco’s Phosphate Buffered Saline (PBS) (ATCC) and collected by scraping. Jurkat T cells were grown in suspension in T-75 flasks (Fisher Scientific, Inc.) and harvested by centrifugation followed by two PBS washes. Replicate plates of cells were prepared for cell counting. Cell counts were obtained from TC20 Automated Cell Counter (Bio-Rad, Hercules, CA, USA).

### Tissue Specimens

Fresh-frozen (FF) tumor specimens (stage IIIC) were obtained from primary (adnexal mass) or metastasis (omentum) of five patients with a primary HGSOC disease site originating at the ovary or fallopian tube. Chemotherapy-naïve and neoadjuvant chemotherapy (NACT) individuals between the age of 40 – 76 years at the time of diagnosis were included (Supplementary Table 1). The study protocol was approved for use under a Western IRB-approved protocol (“An Integrated Molecular Analysis of Endometrial and Ovarian Cancer to Identify and Validate Clinically Informative Biomarkers”) deemed exempt under US Federal regulation 45 CFR 46.102(f). All experimental protocols involving human data in this study were in accordance with the Declaration of Helsinki and informed consent was obtained from all patients.

### Laser Microdissection (LMD)

FF tissue sections (10 µm thickness) were cut by cryostat and placed onto PEN membrane slides (Leica Microsystems, Deer Park, IL, USA). After staining with aqueous hematoxylin and eosin, cell type annotation and counting were performed using the HALO image analysis software (Indica Labs, Albuquerque, NM, USA). Regions of tissue annotated using HALO were exported for LMD (LMD7, Leica Microsystems) as previously described (30). LMD harvested tissue for protein digestion was collected in MircoTubes (Pressure Biosciences, Inc., South Easton, MA, USA) containing 20 µL of 100 mM triethylammonium bicarbonate (TEAB, pH 8.0)/10% acetonitrile, capped and stored at −80 °C until digestion. Tissue for RNA isolation was collected in 300 µL of Buffer RLT with 10% β-mercaptoethanol (QIAGEN Sciences, LLC, Germantown, MD, USA) and stored at −80 °C until isolation.

### Pressure Cycling Technology Trypsin Digestion of Cells and Laser Microdissected Tissues

Cells were transferred to MicroTubes for a final volume of 20 µL of 100 mM TEAB (pH 8.0)/10% acetonitrile. Cell and tissue samples underwent pressure-assisted trypsin digestion employing a barocycler (2320EXT, Pressure BioSciences, Inc.) and a heat-stable form of trypsin (SMART Trypsin, Thermo Fisher Scientific, Inc., Waltham, MA, USA). Peptide digest concentrations were determined using the bicinchoninic acid assay (BCA; Thermo Fisher Scientific, Inc.). Peptides (10 µg for cells and 4 µg for tissue samples) were labeled with isobaric tandem mass tag (TMT) reagents according to the manufacturer’s instructions (TMTpro, Thermo Fisher Scientific, Inc.). Sample multiplexes were reversed-phase fractionated (basic pH) on a 1260 Infinity II offline liquid chromatography system (Agilent Technologies, Inc., Santa Clara, CA, USA) into 96 fractions using a linear gradient of acetonitrile (0.69% min^-1^) followed by concatenation into 36 pooled fractions. Each pooled fraction was resuspended in 25 mM NH_4_HCO_3_ and analyzed by LC-MS/MS.

### RNA Isolation from Cells and Laser Microdissected Tissues

Total RNA was isolated using TRIzol (Thermo Fisher Scientific, Inc.) according to the manufacturer’s instructions and cleaned up with DNase treatment using the RNeasy Mini Kit (QIAGEN Sciences, LLC) for cell samples. RNA from tissue samples were purified using the RNeasy Micro Kit (QIAGEN Sciences, LLC.) per the Purification of Total RNA from Microdissected Cryosections Protocol including on-column DNase digestion. Initial RNA concentrations and 260/280 absorbance ratios were determined using a Nanodrop 2000 Spectrophotometer (Thermo Fisher Scientific, Inc.). Final RNA concentrations were determined using 1 µL of sample and the Qubit RNA HS kit (Thermo Fisher Scientific, Inc.). RNA integrity numbers (RINe) were calculated using the High Sensitivity RNA ScreenTape and Buffers on a Tapestation 4200 (Agilent Technologies, Inc.).

### Generation of *in vitro* Cell Admixtures for LC-MS/MS Analysis

Pooled admixtures (n=13) were generated by combining peptide digests (10 µg total) from OVCAR-3, tumor associated fibroblasts, and Jurkat T cells at pre-defined compositional ratios representing quartile dilutions of cell populations of interest (Supplementary Table 2). Similarly, pooled admixtures (n=17; 4 µg total) were generated by combining peptide digests from LMD enriched populations of tumor, stroma, and lymphocytes (Supplementary Table 3). Cell admixture conditions generated from LMD enriched tissues reflected mainly quartile dilution series, but also included select dilution combining <25% lymphocytes and tumor populations due to limited yields from lymphocyte admixed collections. Digests from multiple patients were combined to make cell type-associated pooled samples. Duplicate digest samples (from cells or tissues) were prepared for models containing a single cell type. The percentage of each cell type in a cell admixture was calculated based on µg of digest.

### Generation of *in vitro* Cell Admixtures for RNA-seq Analysis

RNA pooled admixtures (n=16) were generated by combining RNA samples (20 ng total) from OVCAR-3, tumor associated fibroblasts, and Jurkat T cell at pre-defined compositional ratios (Supplementary Table 4). Similarly, cell type-associated RNA pooled admixtures (n=23) were generated by combining isolated RNA (20 ng total) from LMD enriched populations of tumor, stroma, and lymphocytes (Supplementary Table 5). Cell admixture conditions generated from LMD enriched tissues reflected mainly quartile dilution series, but also included select dilution combining <25% lymphocytes and tumor populations due to limited yields from lymphocyte admixed collections. Isolated RNA from multiple patients were combined to make cell type-associated pools. Duplicate RNA samples (from cells or tissues) were prepared for models containing a single cell type. The percentage of each cell type in a cell admixture was calculated from ng of RNA.

### LC-MS/MS Analysis

Liquid chromatography-tandem mass spectrometry (LC-MS/MS) analyses of TMTpro16 and TMTpro18 multiplexes were performed on a nanoflow high-performance LC system (EASY-nLC 1200, Thermo Fisher Scientific, Inc.) coupled online with an Orbitrap mass spectrometer (Q Exactive HF-X, Thermo Fisher Scientific, Inc.). Samples were loaded on a reversed-phase trap column (Acclaim^TM^ PepMap^TM^ 100 Å, C-18, 20 mm length, nanoViper Trap column, Thermo Fisher Scientific, Inc.) and eluted on a heated (50 °C) reversed-phase analytical column (Acclaim^TM^ PepMap^TM^ RSLC C-18, 2 μm, 100 Å, 75 μm × 500 mm, nanoViper, Thermo Fisher Scientific, Inc.) by developing a linear gradient from 2% mobile phase A (2% acetonitrile, 0.1% formic acid) to 32% mobile phase B (95% acetonitrile, 0.1% formic acid) over 120 min at a constant flow rate of 250 nL/min. Full scan mass spectra (MS) were acquired using a mass range of *m/z* 400-1600, followed by selection of the top 12 most intense molecular ions in each MS scan for high-energy collisional dissociation (HCD). Instrument parameters were as follows: Full MS: AGC, 3 × 10e6; resolution, 60 k; S-Lens RF, 40%; max IT, 45 ms; MS2: AGC, 10e5; resolution, 45 k; max IT, 95 ms; quadrupole isolation, 1.0 *m/z*; isolation offset, 0.2 *m/z*; NCE, 30; fixed first mass, 100; intensity threshold, 2 × 10e5; charge state, 2–4; dynamic exclusion, 20 s, TMT optimization. Global protein-level abundances were generated from peptide spectral matches identified by searching .raw data files against a publicly available, non-redundant human proteome database (http://www.uniprot.org/, SwissProt, Homo sapiens, downloaded 12-01-2017 for cell line and 10-29-2021 for tissue) using Mascot (Matrix Science, v2.6.0), Proteome Discoverer (v2.2.0.388, Thermo Fisher Scientific, Inc.), and in-house tools using identical parameters as previously described (31). Quan correction was applied to all reagent ion abundances using TMTpro16 reagent lot UL296296 or TMTpro18 reagent lots WK334339 and WJ338613.

### Library Preparation and HiSeq Sequencing

RNA library preparation and sequencing were conducted at GENEWIZ, LLC. (South Plainfield, NJ, USA). SMART-Seq v4 Ultra Low Input Kit for Sequencing was used for full-length cDNA synthesis and amplification (Clontech, Mountain View, CA, USA). Nextera XT library (Illumina, Inc., San Diego, CA, USA) was used for sequencing library preparation. Briefly, cDNA was fragmented, and adaptor was added using Transposase, followed by limited-cycle PCR to enrich and add index to the cDNA fragments. The final library was assessed with TapeStation (Agilent Technologies, Inc.). The sequencing libraries were multiplexed and clustered on a flowcell. After clustering, the flowcell was loaded on the Illumina HiSeq instrument according to manufacturer’s instructions. The samples were sequenced using a 2×150 Paired End (PE) configuration. Image analysis and base calling were conducted by the HiSeq Control Software (HCS). Raw sequence data (.bcl files) generated from Illumina HiSeq was converted into fastq files and de-multiplexed using Illumina’s bcl2fastq 2.17 software. One mismatch was allowed for index sequence identification. After investigating the quality of the raw data, sequence reads were trimmed to remove possible adapter sequences and nucleotides with poor quality using Trimmomatic v.0.36. The trimmed reads were mapped to the *Sus scrofa* reference genome available on ENSEMBL using the STAR aligner v.2.5.2b. The STAR aligner is a splice aligner that detects splice junctions and incorporates them to help align the entire read sequences. BAM files were generated as a result of this step. Unique gene hit counts were calculated by using feature counts from the Subread package v.1.5.2. Only unique reads that fell within exon regions were counted. GRCh38/hg38 was used as the human reference genome. Count level data was VST normalized by DESeq2 (ver 1.24.0).

### Analysis of Deconvolution Tools and Prognostic Molecular Subtypes

Proteomic data from admixed samples comprised of 33.3% OVCAR-3/tumor, 33.3% fibroblast/stroma, and 33.3% Jurkat/lymphocytes were used to normalize protein abundances. Global proteome data was visualized by principal component analyses (PCA) using the top 100 most variable proteins or transcripts by mean absolute deviation (MAD) using ggplot2 (version 3.4.1) in R (version 4.2.2) (32). Gene names in the proteomic and RNA-seq datasets were harmonized with any gene synonyms used in the Consensus^TME^ and ESTIMATE tools prior to their utilization. RNA-seq data was filtered by excluding features without HUGO Gene Nomenclature Committee (HGNC) gene symbols, gene symbols that began with “LOC”, or zero variant entries. RNA-seq data was gene-wise, z-score scaled and subsetted to genes co-quantified in proteomics for use with ESTIMATE and Consensus^TME^. Pearson correlation analysis was performed between ESTIMATE or Consensus^TME^ and percent cell type (quartile cell percentages). Transcriptomic data from the admixed samples comprised of 33.3% OVCAR-3/tumor, 33.3% fibroblast/stroma, and 33.3% Jurkat/lymphocytes were excluded from the correlation analysis (outliers based on ESTIMATE and Consensus^TME^ scores). Spearman correlation was performed on transcriptomic data and proteomic data from tissue admixture models and heatmap of quartile percentage tissue models were generated using ComplexHeatmap (version 2.14.0) (33). RNA-seq data (quartile cell percentages) from HGSOC tissue cell admixture was used for subtype classification with consensusOV (version 1.20.0) (10). Proteomic data from tissue admixture models were assessed by Decomprolute (21).

### Cell Type Protein Signature Prioritization and Verification

Differential analysis was performed using limma (34) to identify proteins uniquely elevated within cell populations of interest enriched from tissue using LMD. Protein features of interest passed an adjusted *p value* < 0.05 comparing tumor with stroma/immune collections, immune with tumor/stroma collections and stroma protein features passed a *p value* < 0.05 and as well as a fold-change (FC) cutoff ± 1.5. Protein abundance data (Supplementary Table 6) was then partitioned into three sets of training (n=10) and testing (n=6) samples stratified by tissue composition using the ‘train_test_split’ function in scikit-learn (version 1.2.1). Recursive feature elimination (RFE) on the respective training data was used to select each protein signature that holds the most predictive power for support vector regression models (SVR) with linear kernels. Each SVR model’s target was the percent composition of the respective tissue type. Grid search was utilized to find optimal hyperparameters before performing RFE for each signature. The ranks provided by RFE for each protein signature were then used to re-train SVR models and assess their performance on their respective testing data. The SVM model with at least 15 features and had the lowest mean squared error (MSE) to the test data was chosen, and its features were determined as the protein cell type signatures. Models were trained in python (version 3.9.16) using the scikit-learn package (version 1.2.1) (35). Upset plots were generated using ComplexUpset (version 1.3.5) (36). Transcript-protein Spearman correlations were calculated from matched tissue admixture model data for cell type-associated protein signatures. Assignment of admixed samples with prognostic HGSOC molecular subtypes (12) was performed using consensusOV (ver 1.18.0) (10), where a total of 575 unique Entrez gene IDs (kindly provided by the author) were identified from 635 selected probe sets. Correlation plots were generated with seaborn version 0.11.2. Mann-Whitney Wilcoxon Test was used to assess the statistical significance between groups. Protein signatures were validated using proteomic abundance data from enriched and bulk HGSOC (n = 9) (7) and bulk HGSOC tissues (n = 169) by performing ssGSEA with the newly derived gene sets (37, 38). ssGSEA was performed in R (version 3.6.0) using the GSVA library (version 1.34.0) (38). The top 25% most variably abundant proteins by standard deviation from bulk HGSOC tissues (n = 169) (6) was visualized by principal component analyses (PCA) and overlaid with stroma, immune, or tumor ssGSEA scores. Package scikit-learn (version 1.2.1) was used for performing PCA, while packages seaborn (version 0.11.2), matplotlib (version 3.7.1), and scipy (version 1.10.0) were used for plotting.

## Supplementary Tables

Supplementary Table 1: Patient cohort and cell types collected from each tissue sample.

Supplementary Table 2: Cell admixtures from cell lines for proteomic analysis

Supplementary Table 3: Cell admixtures from HGSOC LMD enriched tissues for proteomic analysis

Supplementary Table 4: Cell admixtures from cell lines for RNA-seq analysis

Supplementary Table 5: Cell admixtures from HGSOC LMD enriched tissues for RNA-seq analysis

Supplementary Table 6: Proteomic data from HGSOC tissue admixtures

Supplementary Table 7: Proteomic data from cell line admixtures

Supplementary Table 8: RNA-seq data from HGSOC tissue admixtures

Supplementary Table 9: RNA-seq data from cell lines admixtures

Supplementary Table 10: Protein and transcript Spearman correlation in admixtures

Supplementary Table 11: ESTIMATE scores across cell line admixtures (proteome data)

Supplementary Table 12: ESTIMATE scores across HGSOC tissue admixtures (proteome data)

Supplementary Table 13: ESTIMATE scores across cell line admixtures (transcriptome data)

Supplementary Table 14: ESTIMATE scores across HGSOC tissue admixtures (transcriptome data)

Supplementary Table 15: Consensus^TME^ scores across cell line admixtures (proteome data)

Supplementary Table 16: Consensus^TME^ scores across HGSOC tissue admixtures (proteomic data)

Supplementary Table 17: Consensus^TME^ scores across cell line admixtures (transcriptome data)

Supplementary Table 18: Consensus^TME^ scores across HGSOC tissue admixtures (transcriptome data)

Supplementary Table 19: Pearson correlation analysis of ESTIMATE and Consensus^TME^ scores with percent cell type

Supplementary Table 20: Decomprolute scores across HGSOC tissue admixtures (proteome data)

Supplementary Table 21: Protein cell type signatures and ranking

Supplementary Table 22: Transcript and protein abundance correlation of co-identified signatures

## Acknowledgements

The authors would like to acknowledge Sakiyah TaQee, Persus Akowuah, Jeremy Loffredo, Glenn Gist, Salma Eltahir, Sasha Makohon-Moore, and Dr. Paulette Fauceglia for their contributions to histopathology assessment and sample preparation, and informatics data analysis. We would also like to acknowledge the patients and families who helped to make this work possible.

## Authors’ contributions

N.W.B. and T.P.C. contributed to conception, experimental design, data analysis and manuscript composition. P.N.T. performed experiments, data analysis and manuscript composition. T.A. and J.S. contributed to bioinformatics and data analysis. J.O., V.O., F.S.P., M.E., A.C., B.L.H., K.A.C., D.M., A.L.H., T.L., P.-K.R.-K., N.T.P., K.M.D. and G.L.M. provided sample preparation, data generation and manuscript review.

## Ethics approval and consent to participate

The study protocol was approved under a Western IRB-approved protocol “An Integrated Molecular Analysis of Endometrial and Ovarian Cancer to Identify and Validate Clinically Informative Biomarkers” deemed exempt under US Federal regulation 45 CFR 46. 102 (f) and informed consent was obtained from all patients involved in the study protocol. All experimental protocols involving human data was performed in accordance with the Declaration of Helsinki.

## Data availability

The mass spectrometry proteomics data have been deposited to the ProteomeXchange Consortium (http://proteomecentral.proteomexchange.org) via the PRIDE (39) partner repository with the dataset identifier PXD044157. RNA-seq and proteomic data are included in the Supplementary tables S8 – S11. The protein signature assessment tool is available at: https://lmdomics.org/ProteoMixture/.

## Disclaimer

The contents of this publication are the sole responsibility of the authors and do not reflect the views, opinions or policies of Uniformed Services University of the Health Sciences, the Henry M. Jackson Foundation for the Advancement of Military Medicine, Inc, the Department of Defense or the Departments of the Army, Navy or Airforce. Mention of trade names, commercial products or organizations do not imply endorsement by the U.S. Government.

---

**Supplementary Figure 1:**
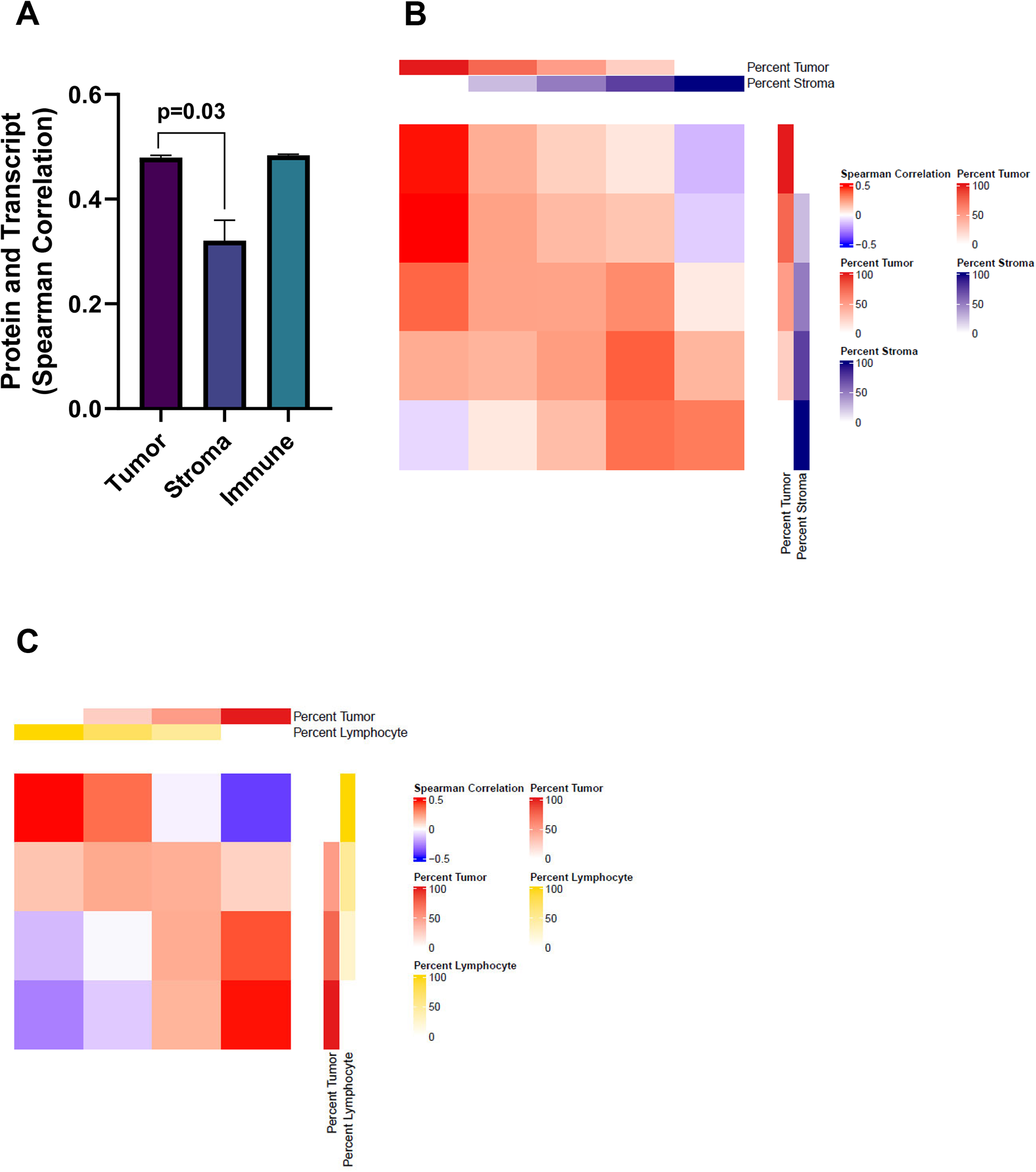
A: Comparison of protein and transcript correlation with tumor, stroma, and immune cell populations enriched from HGSOC tissues. Heatmaps depicting Spearman correlations between tissue admixture samples using proteins and transcripts co-quantified from quartile tumor/stroma (B) and tumor/lymphocyte (C) admixture models.

**Supplementary Figure 2:**
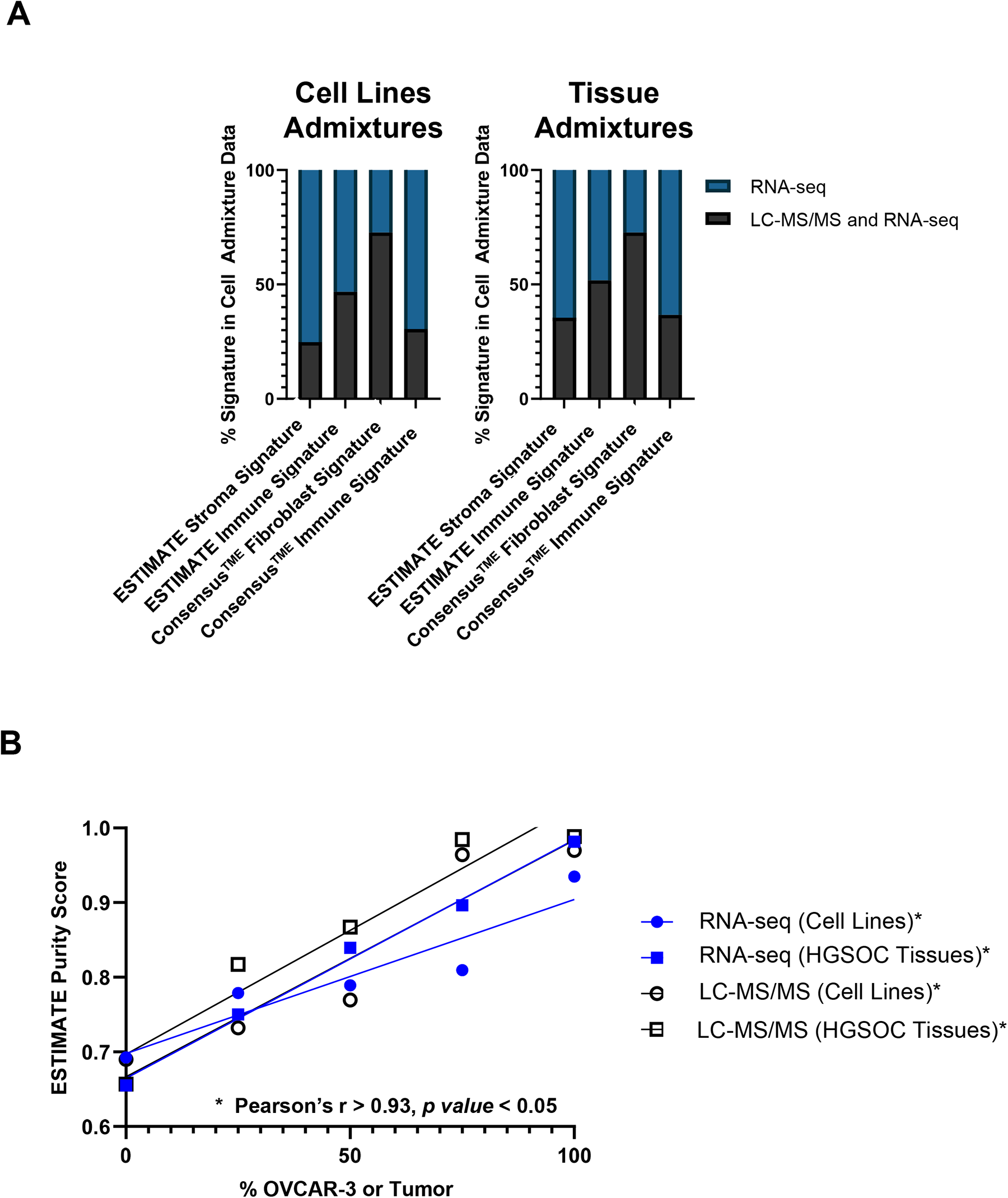
A: Deconvolution tool gene signatures co-quantified in cell admixture proteome and transcriptome data in cell line and tissue collections for ESTIMATE stromal and immune as well as Consensus^TME^ fibroblast and immune. B: Correlation analysis of transcript (RNA-seq) and protein (LC-MS/MS) level scores for quartile dilutions from cell lines and tissue collections corresponding to ESTIMATE tumor purity scores.

**Supplementary Figure 3:**
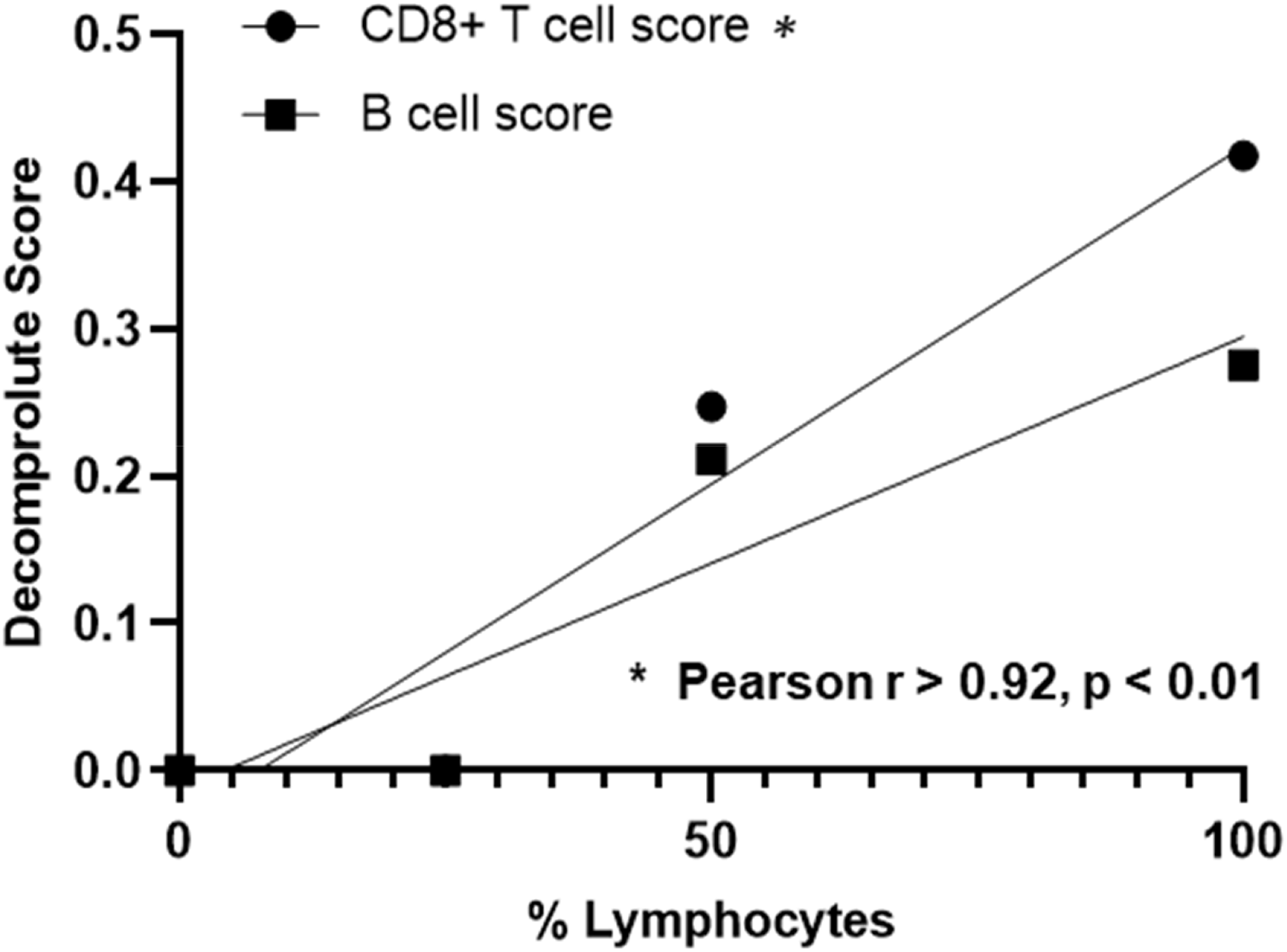
Decomprolute CD8+ T cell and B cell scores across HGSOC tissue admixture models from tumor/ lymphocyte admixed collections using protein-level data.

**Supplementary Figure 4:**
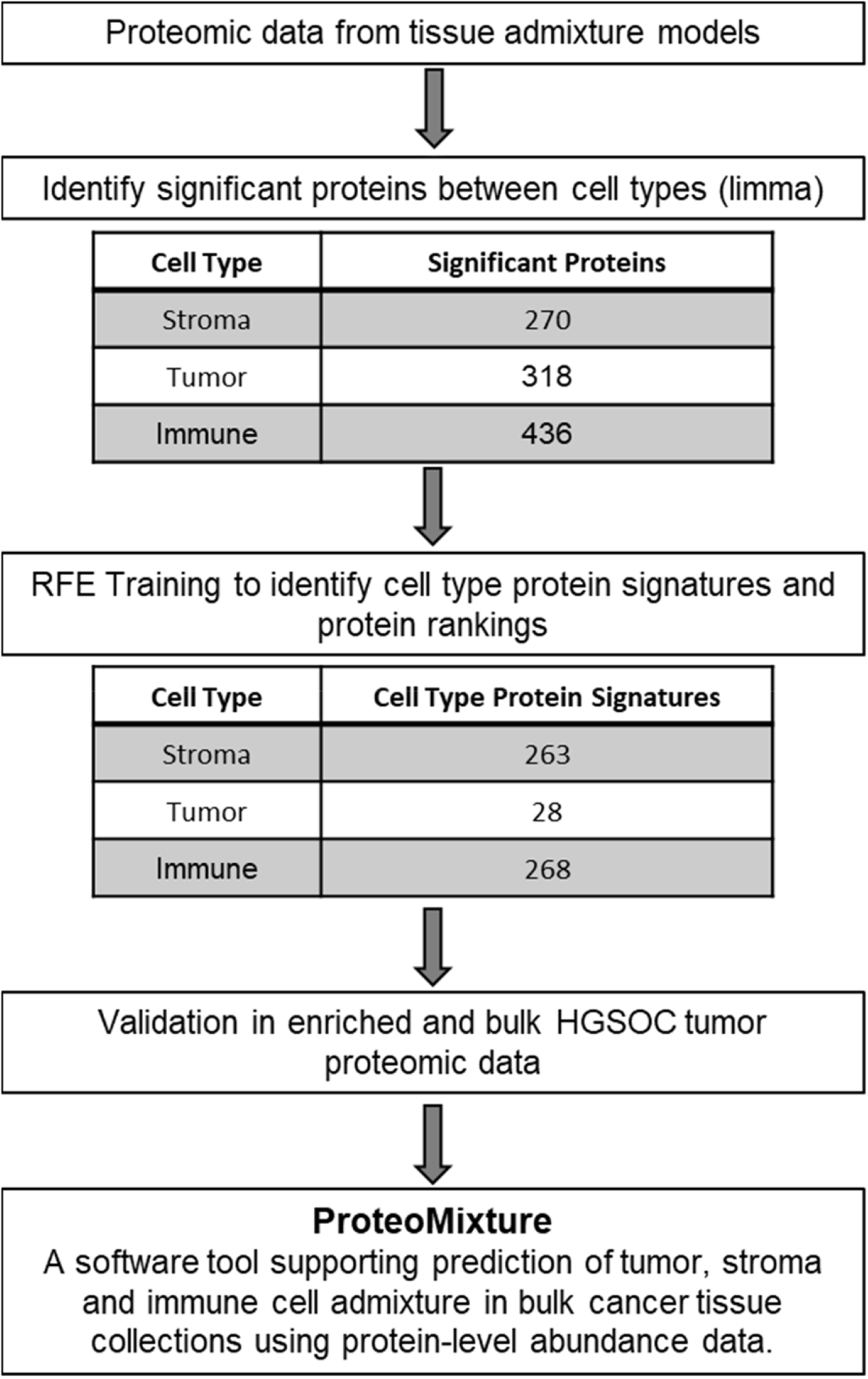
Strategy for prioritization, optimization, and analysis of proteomic cell type signatures.

**Supplementary Figure 5:**
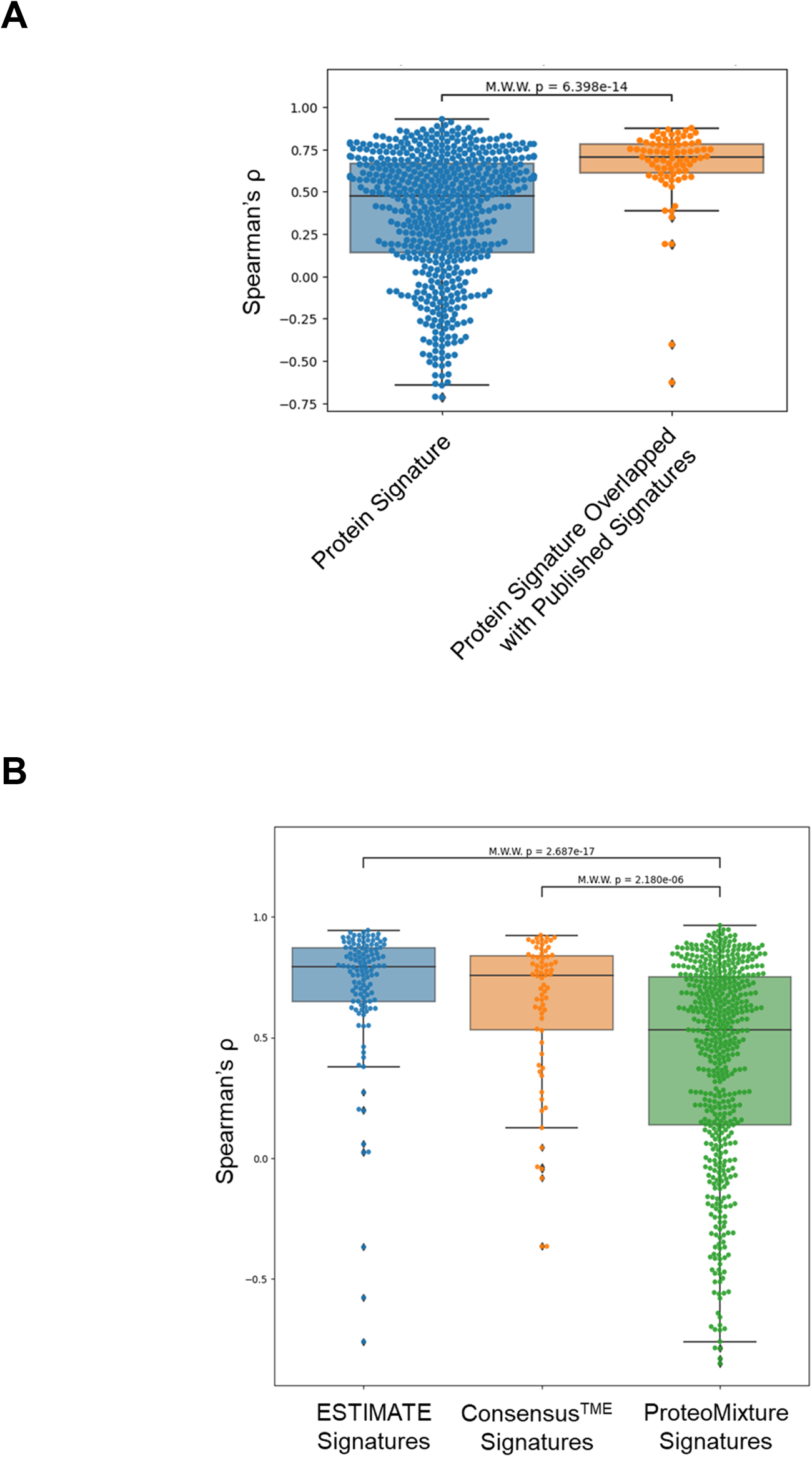
A: Protein and transcript correlation of all protein signatures and protein signatures overlapped with previously published signatures. B: Spearman correlation of protein and transcript from deconvolution tools identified in HGSOC cell admixture dataset. Abbreviation: Mann-Whitney-Wilcoxon (M.W.W.).

**Supplementary Figure 6:**
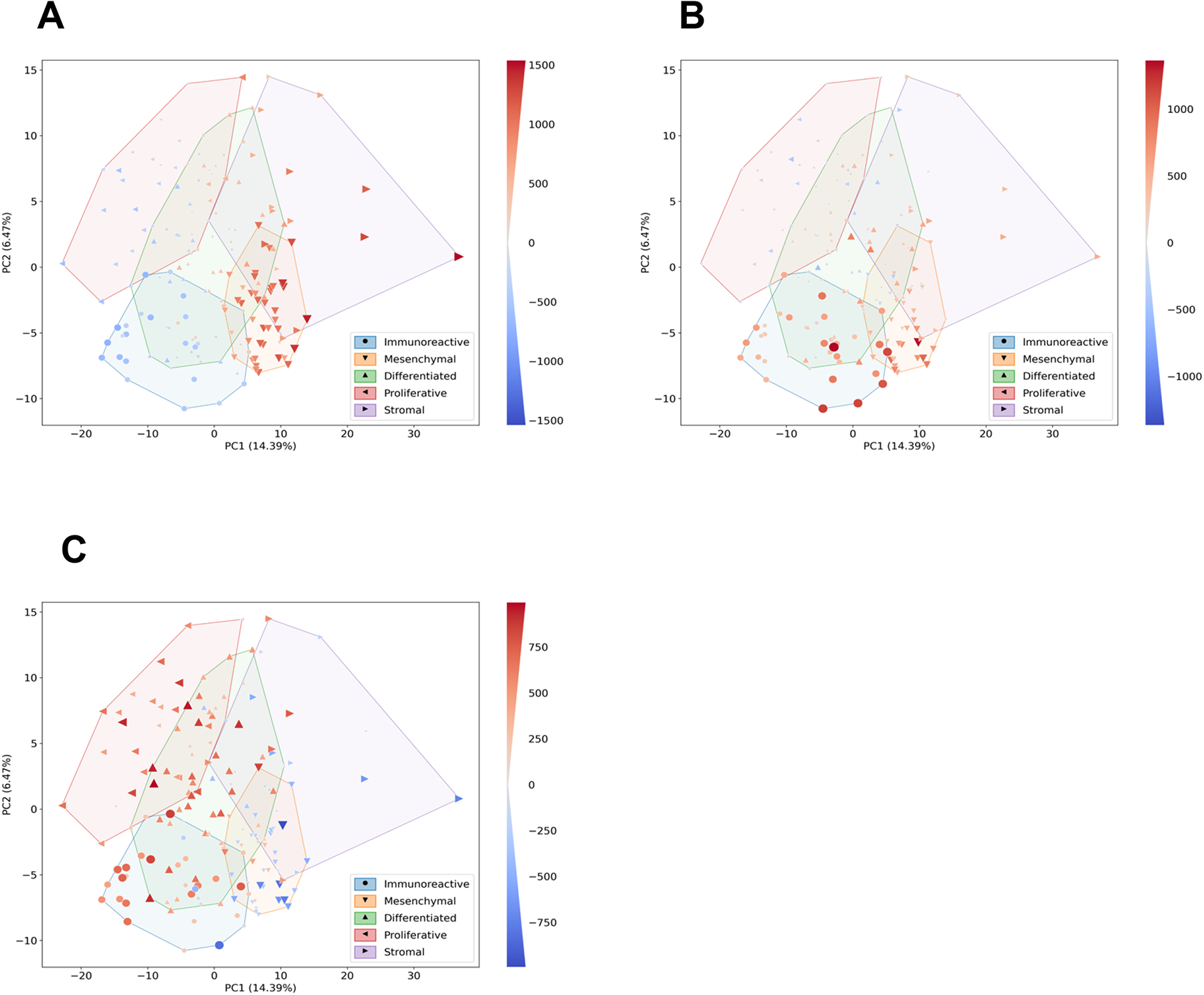
A: PCA of top twenty-five percent most variable proteins by median absolute deviation (MAD) from bulk HGSOC tissues overlaid with ProteoMixture stroma score. B: PCA of top twenty-five percent of proteins by MAD from bulk HGSOC tissues overlayed with immune score. C. PCA of top twenty-five percent of proteins by MAD from bulk HGSOC tissues overlayed with tumor score.

